# HIHISIV: a database of gene expression in HIV and SIV host immune response

**DOI:** 10.1101/2023.11.25.568664

**Authors:** Raquel Lopes Costa, Luiz Gadelha, Mirela D’arc, Marcelo Ribeiro-Alves, David L Robertson, Jean-Marc Schwartz, Marcelo A. Soares, Fábio Porto

## Abstract

In the battle of the host against lentiviral pathogenesis, the immune response is crucial. However, several questions remain unanswered about the interaction with different viruses and their influence on disease progression. The simian immunodeficiency virus (SIV) infecting nonhuman primates (NHP) is widely used as a model for the study of the human immunodeficiency virus (HIV). Both because they are evolutionarily linked and because they share physiological and anatomical similarities that are largely explored to understand the disease progression. The HIHISIV database was developed to support researchers to integrate and evaluate the large number of transcriptional data associated with the presence/absence of the pathogen (SIV or HIV) and the host response (NHP and human). The datasets are composed of microarray and RNA-Seq gene expression data that were selected, curated, analyzed, enriched, and stored in a relational database. Six query templates comprise the main data analysis functions and the resulting information can be downloaded. The HIHISIV database provides accurate resources for browsing and visualizing results and for more robust analyses of pre-existing data in transcriptome repositories.

**Database URL:** https://hihisiv.github.io

## Background

The evolutionary history of Human Immunodeficiency Virus type 1 (HIV-1) is closely linked with *Simian Immunodeficiency Virus* (SIV), with four cross-species introductions described from chimpanzees (HIV-1 M and N) and gorillas (HIV-1 O and P) into humans. For years, the general thought was that all African monkey species do not develop immunodeficiency syndrome when infected with SIVs, namely natural hosts, similar to the human cases of non-progressive and controller patients (1) (2) (3) (4) (5) (6). However, little is known about the *in vivo* pathogenicity of SIV because it is very difficult to monitor infected apes in the wild. Only captive data were initially analyzed and it was assumed that all African monkeys develop a non-pathogenic phenotype, such as sooty mangabeys and African green monkeys that are widely studied in captivity. Also in captivity, Macaque species, endemic to Asia, can be experimentally infected with virus strains from sooty mangabeys and progress to an AIDS-like phase. The clinical hallmarks are similar to the common human disease, namely non-natural hosts and progressive patients, respectively (7) (8). Nevertheless, epidemiological surveys in free-ranging chimpanzees showed that SIVcpz has a substantial negative impact on the health, reproduction and lifespan.

One of the major challenges in these viruses is understanding the complex interplay between the host’s immune system, which may involve the depletion of TCD4+ lymphocytes. In the last decades, the advances in high-throughput technologies enabled analyzing a large number of transcripts simultaneously and inferring subsets of these transcripts that are associated with the same biological response (9). Although the amount of data is constantly increasing in transcriptome repositories, such as GEO (Gene Expression Omnibus), ArrayExpress, and SRA (Sequence Read Archive), the data are not processed, standardized, and integrated.

HIHISIV is a database of host immune response in expression profiles of SIV and HIV and was developed to support the researchers in identifying molecular signatures, genes co-expressed, experimental design, and hosts.

The database is composed of microarray and RNA-Seq gene expression data of viral infection by SIV and HIV hosts retrieved from the GEO repository. The datasets were manually curated and reanalyzed using standardized methods and the metadata and results were annotated and enriched with ontology terms. The HIHISIV database is currently in version 2.0 and includes 63 transcriptome experiments stored in a relational database. Furthermore, HIHISIV has a user-friendly web interface where co-expression networks can be visualized as graph and tabular data views and the dataset can be downloaded for further analysis.

## Construction and Content

### Datasets & Data Analysis

#### Dataset selection

The initial datasets consists of microarray (1-color mRNA) and RNA-Seq (mRNA) gene expression data related to viral infections by SIV or HIV-1 in the host, retrieved from the GEO from the National Center for Biotechnology Information (NCBI). We compiled the initial database by selecting searches to ‘SIV’ or ‘HIV-1’ projects. Each dataset were manually examined to identify and separate the groups for contrast (reference *versus* test). Additionally, we established exclusion criteria for GEO projects or samples, and the following criteria were adopted for removing projects or samples:

i) Lack of sample information preventing the differentiation of groups for comparison (e.g., host type); ii) Projects or samples related to antiretroviral therapy (ART), vaccination, monoclonal antibodies, or cell cultures; iii) Insufficient number of samples (less than 3 per group).

As a proof of concept to create the pipeline and compose the conceptual model, we selected one short-read RNA-Seq project (GEO id: GSE119234) and applied the same exclusion criteria mentioned before.

During the curation process, we encountered various terminologies that refer to the same entity. To ensure standardization across different datasets and establish consistency, we have adopted the following nomenclature:

- natural-host: refers to natural host primates such as African Green Monkeys (*Chlorocebus aethiops*) and Sooty mangabeys (*Cercocebus atys*);
- non-natural-host: refers to non-natural host primates such as Rhesus monkey (*Macaca mulatta*);
- non-progressor: refers to a human that has not progressed to immunodeficiency.
- uninfected (EFO_0001460): Uninfected class is a disposition in which the bearer is not known to be affected by a disease withtin the context of a study.
- acute infection (IDO:0000627): An infectious disorder that is the physical basis for an unfolding acute infectious disease course.
- chronic infection (IDO:0000628): An infection that persists for an extended period of time.

In each project, the data was compared pairwise, and each comparison was named as an *experiment*. In some cases, these projects resulted in more than one experiment. For example, the project GSE7157 worked with non-natural species during three different phases of the infection (uninfected, acute and chronic infections). This project was divided into three different pairwise experiments (uninfected *versus* acute: GSE7157_d1; uninfected *versus* chronic: GSE7157_d2, and acute *versus* chronic: GSE7157_d3). The complete list of experiments is shown in Experiments list in the database web page.

#### Metadata and Ontologies

To enrich the experimental information with ontology terms, we employed the GEOquery library, for retrieving metadata (including phenodata) from GEO datasets. The extracted metadata consisted of semi-structured information, including essential details in the experiments as title, summary, and overall design. We used SpaCy model, a natural language processing (NLP) tool, to identify automatically the key elements from these metadata (19). After that, the spurious or irrelevant terms was carefully filtered and the remain terms were mapped to their corresponding biological ontologies using rols library (20) to access and query the Ontology Lookup Service (OLS) improving the interoperability and integration of HIHISIV database with other sources.

#### Microarray and RNA-Seq data analysis

Due to the heterogeneity of experiments as conducted by different research teams, assay platforms, virus strains, and tissues, the analyzes were conducted separately through the pipeline delineated in a dataflow shown in Figure 1. The list of experiments with the correspondent normalization matrix and phenodata are available in the HIHISIV database.

**Figure 1:**
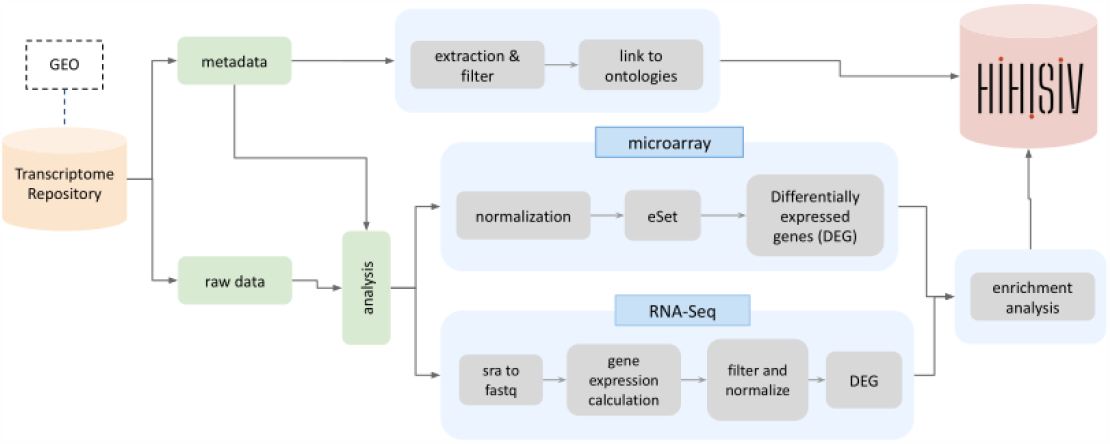
Dataflow used to generate data for HIHISIV showing metadata and raw data input, their RNA-Seq, microarray, and enrichment analyses, the metadata extraction of relevant entities, and insertion of all results in the database.

All microarray data (1-color mRNA) were downloaded from the Affymetrix platform. The datasets were normalized using the MAS5 normalization method implemented in *Affy* R package (version 1.78.2) (13) resulting in a normalized matrix with the probes. For each experiment, a pair group was compared. The first condition was considered as the reference and the second condition as the test, for instance, uninfected (reference) *versus* acute infection (test).

Finally, the detection of the differentially expressed genes (DEGs) was based on normalized datasets by the fitting of the gene-wise linear model (for each probe) followed by moderated t-tests implemented in the *limma* package in R (version 3.56.2) (15). We calculated the up and down-regulated genes between the sample pairs through the adjusted p-value and the log fold-change measures. We kept the complete results of the analysis in the database and probes mapped to more than one entrez_gene_id were inserted as separate tuples in the database. The orthologous genes in *Macaca mulatta* and *Homo sapiens* were mapped using *bioRmat* R library (11).

To establish a standardized pipeline a RNA-Seq (mRNA) dataset was collected from the SRA repository (SRS from NextSeq 550 *Homo sapiens*). Firstly, the data were transformed to Fastq using sratoolkit (version 2.10.5) and the quality of reads was checked using FastQC. RSEM *rsem-calculate-expression* (version 1.3) was used to align the transcriptome dataset in the genome reference (UCSC hg19) and after the mapping, the DEGs were evaluated using the library *limma* (limma-voom). The annotation was conducted by the R libraries *TxDBHsapiens, HomoSapiens*, and *GenomicFeatures* (version 1.52.1) (16). The gene identifiers (‘ucsc_id’) were kept.

#### Enrichment analysis

We conducted enrichment analysis using the hypergeometric test implemented in the GOStats package (21) to identify relevant Gene Ontology (GO) terms associated with biological processes (12). The gene subset was defined by the set of differentially expressed (DE) genes obtained from pairwise comparisons between a test and a reference condition. To mitigate potential bias, as demonstrated by Timmons, Szkop & Gallagher (2015) (22), we applied a more restrictive cut-off (p-value < 0.001) to reduce detection bias.

### Database, Design, and Description

To reproduce the concepts and relationships among entities such as genes, ontologies and experiments, we represent this scenario using a conceptual model (Figure 2) and the main entities are described below.

**Figure 2:**
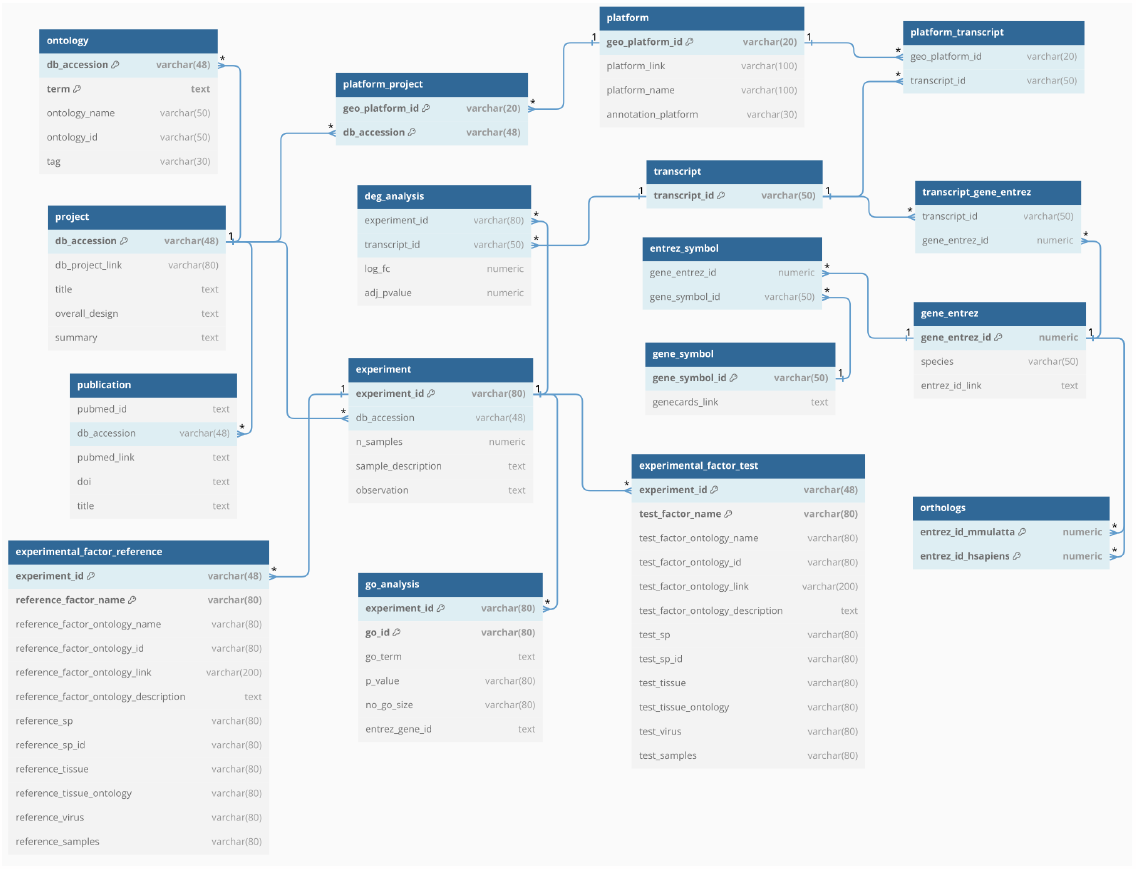
The HIHISIV conceptual model representing the database entities and their relationships. One of the main entities is ‘experiment’, which describes the experiments derived from the project (one or more).

* **project**: describes information about the original project from the GEO. Contains db_accession (key), db_project_link, title, overall_design, and summary.
* **publication**: contains information about publication(s) related to the project. The attributes are pubmed_id, pubmed_link, doi, and title.
* **platform:** describes the kind of high-throughput gene expression platform used - RNA-Seq or microarray platforms. Contains information about GEO platform id (geo_platform_id), platform_link, platform_name, and annotation_platform.
* **experiment**: describes the experiments derived from the project (one or more). This entity contains information about the experiment_id (key), number of samples (n_samples), sample_description and observation.
* **experimental_factor_test**: contains attributes related to the group test such as the name factor used as comparisons (test_factor_name), the ontology name associated with the test_factor_name (test_factor_ontology_name), the ontology id and link, the species in the test group (test_sp), the tissue (test_tissue) and the ontology associated with this tissue (test_tissue_ontology), the virus lineage (in case of non-human) and the samples associated with this group.
* **experimental_reference_test**: similar to the entity experimental_factor_test but containing information on the reference group.
* **transcript**: it has a unique attribute describing the probe_id (key) related to the platform.
* **gene_symbol:** contains information about the gene and the attributes are: entrez_gene_id (key), gene_symbol (official HUGO name), link_ncbi, gene_name_desc, and chromosome.
* **deg_analysis**: this entity contains the result of the DEG analysis by RNA-seq or microarray platforms. The attributes are experiment_id, transcript_id, log_fc values and adj_pvalue.
* **go_analysis**: this entity contains the results of the enrichment analysis. The attributes provided are gene ontology id (go_id), go_term, p-value, number of gene in the ontology id (no_go_size), and entrez_gene_id which represents the entrez_gene_id associated with this go_id in the result analysis.
* **orthologs**: this entity mapp the entrez_gene_id in *H. sapiens* to entrez_gene_id in *M. mulatta* (that is the common microarray platform used in the host of SIV experiments).

### System Architecture, Implementation, and Access

The HIHISIV database was designed and implemented as a multiple-component framework, as represented in Figure 3. The database was implemented and instantiated in a PostgreSQL (version 15.3) server following the same conceptual model described previously (Figure 2). The web application provides a user-friendly interface implemented in the *Streamlit* (version 1.25.0) Python library with predefined parameterizable queries to be executed in the database. It enables the analysis of gene expression levels in HIV-1 and SIV hosts. The results are displayed in tables or networks, and all the data can be downloaded by the users. Other Python libraries used in the implementation include *Psycopg2* (version 2.9.5), for connecting to the PostgreSQL database and executing queries; *Pandas* (version 1.5.3) for manipulating data frames; and *NetworkX* (version 3.1) for building networks.

**Figure 3:**
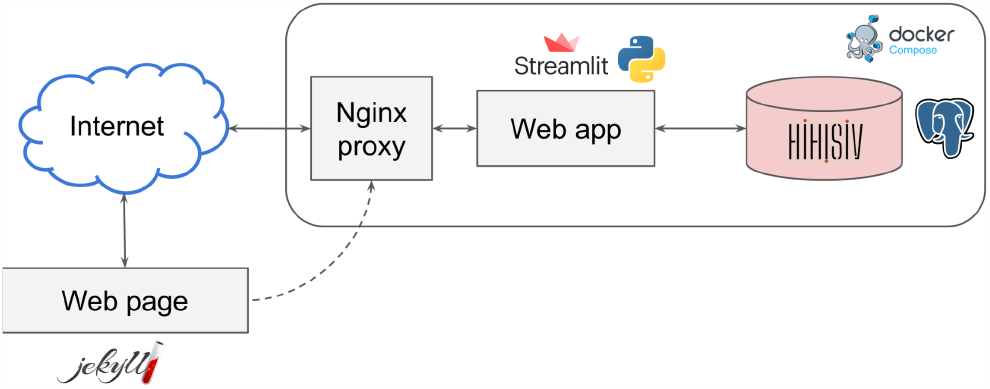
The HIHISIV framework system architecture.

The static web page contains a description of how the database was built and a link to the source data that were used. The web page was written using the Jekyll framework. Finally, the Nginx proxy was used to route the requests from the Internet to the web app. For each of these components, a docker container was built and docker-compose was used to connect and deploy them. The system is hosted at the Data Extreme Laboratory (DEXL) of the National Laboratory for Scientific Computing (LNCC) in Brazil.

### Support for the FAIR Principles

The FAIR principles (17) are considered a gold standard for research data management. In this work, we follow best practices for improving the support for them in the HIHISIV database. For improving findability, basic metadata using schema.org (18) was added to the headers of the HIHISIV web page, allowing for indexing by search engines and dataset repositories. HIHISIV data and metadata are openly accessible through its URL (https://hihisiv.github.io) using the HTTP protocol. When applicable, terms used both in the database schema and in its web interface follow Gene Ontology (12) and other ontologies such as EFO (The Experimental Factor Ontology), UBERON (Uber-anatomy ontology), PATO (the Phenotype And Trait Ontology), MONDO (Mondo Disease Ontology) and NCIT (The National Cancer Institute Thesaurus) improving interoperability. For better reusability, the database contains references to the original data used in the analyses. The source code for the analysis workflows, the database schema and initialization scripts, and the web application is available on Github (https://github.com/quelopes/HIHISIV). This article describes v2.0 of the database, which is archived on Zenodo under DOI: 10.5281/zenodo.7093185.

## Utility and Discussion

### Querying in the User Interface

HIHISIV can be queried through a web interface in which the user can configure and adjust the parameters of a query. There are six query interfaces with major analytic functions, which are ‘*Gene*’, ‘*Transcripts*’, ‘*Biological Process (GO)*’, ‘*Ontology terms*’, ‘*Single gene co-expression network*’, and ‘*Gene set co-expression network*’. The resulting information can be exported in comma-separated values (CSV) file for further data manipulation. Public access and documentation are freely available through the database web page. Next, we provide a set of illustrative examples in a biological context to demonstrate how researcher can utilize HIHISIV to explore relevant datasets, identify key genes associated with immune responses, analyze co-expression networks, and elucidate relevant biological processes related to SIV and HIV-1 infections.

#### Gene

In this mode, the user can select the conditions under which a gene of interest shows differential expression. As an example, Figure 4 presents the results for the gene USP18, (Ubiquitin specific peptidase 18) that belongs to the ubiquitin-specific proteases (UBP). The *USP18* gene and its protein product, *USP18*, are known to be involved in the innate immune response to viral infections, particularly in response to type I interferons, which are cytokines produced by the body during viral infections (23). The applied thresholds for differential expression analysis were adjusted p-value (adj_pvalue) <= 0.05 and |log-FC| > 1. The results show this gene was DE in seven experiments, for example in the comparison reshus uninfected *versus* acute infection (GSE16147_d5, GSE17626_d1, GSE61766_d1), humans uninfected *versus* acute infection (GSE6740_d18) and human non-progressor *versus* acute infection (GSE6740_d21).

**Figure 4:**
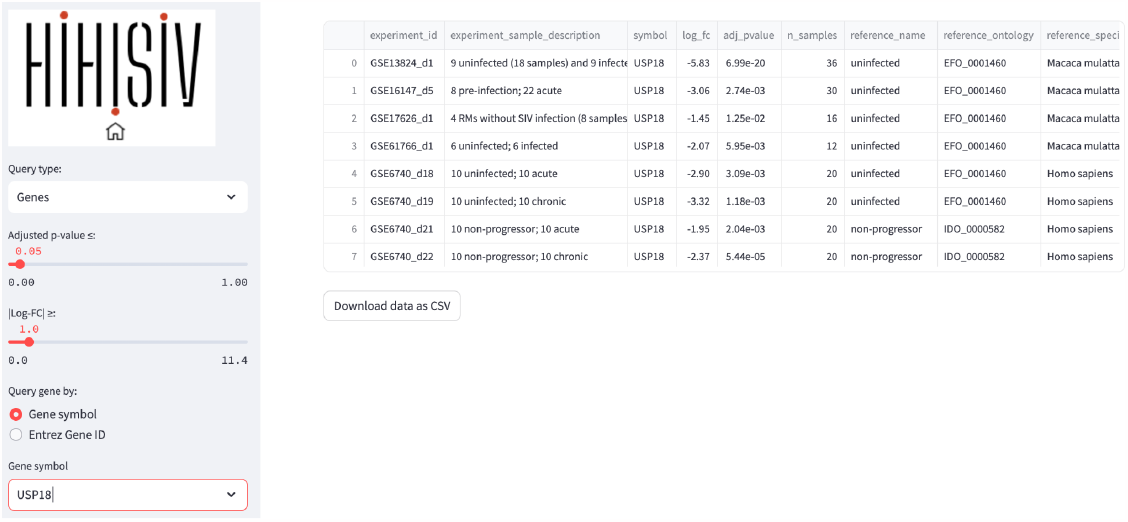
‘Gene’ query example for the gene *USP18*, showing the conditions under which the gene was selected (adj_pvalue <= 0.05 and |log-FC|) > 1).

#### Transcript

This mode is similar to the previous one, but instead of the querying by gene symbol or entrez_gene_id, the user may be interested in a specific probe or transcript id. In the example shown in Figure 5, the probe id ‘*Mmu*.*STS*.*4748*.*a*.*s1_at*’ is selected based on the criteria of adj_pvalue <= 0.05 and |log-FC| > 1. As this probe id is a specific to microarray platforms in *M. mulatta* (GPL3535 - Affymetrix Rhesus Macaque Genome Array) the results show only experiments that used this platform.

**Figure 5:**
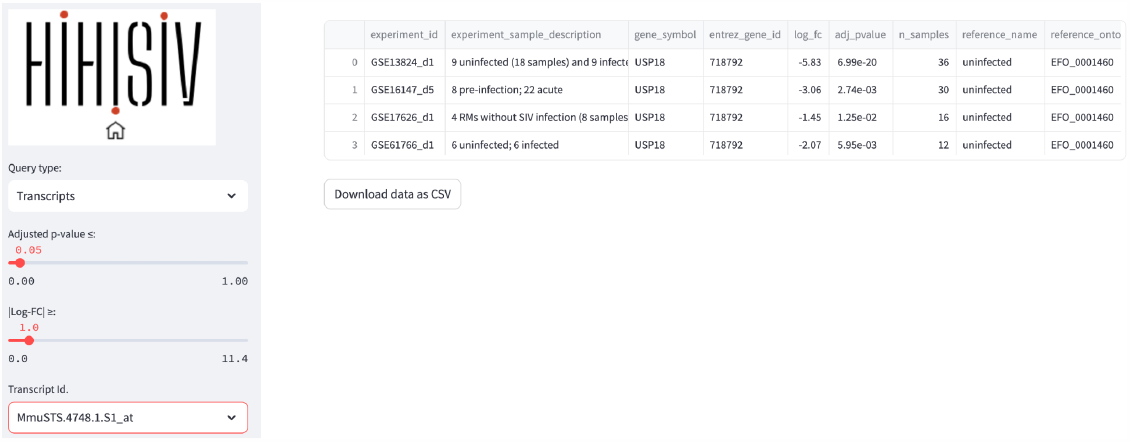
‘Transcripts’ query example shows the result for the probe id ‘Mmu.6048.1.S1_s_at’ (adj_pvalue <= 0.05 and |log-FC| > 1).

#### Gene Ontology (Biological Process)

This mode presents the results of the enrichment for GO terms related to BP for each comparison derived from the DEG analysis. By utilizing this mode, researchers can identify experiments that exhibit enrichment in specific BP terms of interest. For instance, in Figure 6, the BP term ‘*cellular response to type II interferon*’ (GO:0071346) was found enriched in the experiments GSE6740_d1 (CD4+ *versus* CD8+ in acute infection), GSE6740_d2 (CD4+ *versus* CD8+ in chronic infection) and GSE13824_d1 (uninfected *versus* infected). Additionally, this mode provides information about the genes that were enriched within this specific ontology. To ensure statistical robustness, only the results with a significant p-value < 0.001 are displayed in the database.

**Figure 6:**
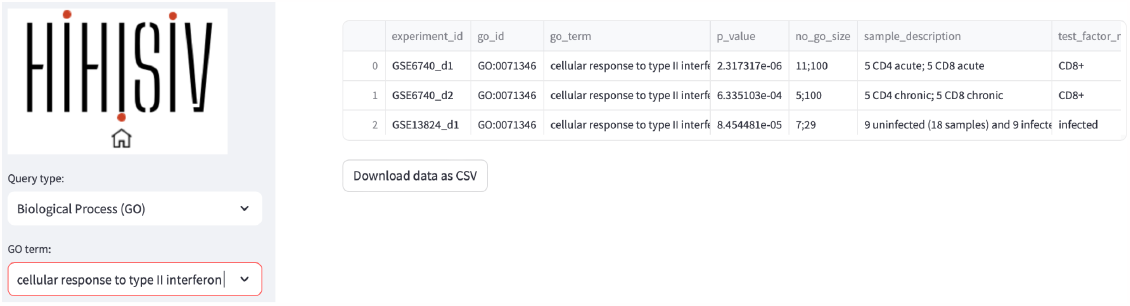
An example of a ‘Gene Ontology’ query shows the result for the biological process domain for the GO term ‘cellular response to type II interferon’.

#### Ontology terms

By integrating ontology terms with the experimental information, our approach provided a comprehensive understanding of the underlying biological contexts and enriched the dataset’s metadata with valuable semantic annotations. In the ‘*Ontology term*’ mode the example in Figure 7 shows the experiment associated with the term ‘*Encephalitis*’ (project GSE13824).

**Figure 7:**
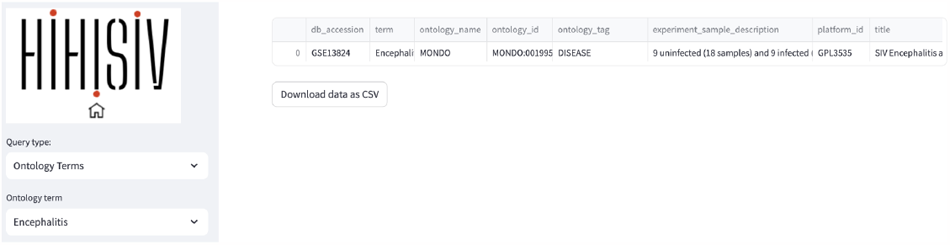
‘Ontology term’ mode shown the project (GSE13824) associated with the term ‘Encephalitis’.

#### Single gene co-expression network

This query mode ‘*Single gene co-expression network*’ represents the genes that were co-expressed with a selected target gene. In the example shown in Figure 8, we are using the same gene as in the ‘Gene’ mode, *USP18* (adj_pvalue <= 0.05 and |log-FC| > 1). The result displays a set of genes that are co-expressed with *USP18* in different experiments (e.g., *IFIT1, PCLAF, IFI35, ISG15* and *OAS2*). The thickness of the connections between genes depends on the number of co-expressed experiments that the target gene has with the resulting gene. For instance, *IFIT1* (interferon induced protein with tetratricopeptide repeats 1) is a gene that encodes a protein that may inhibit viral replication and translational initiation (provided by RefSeq, Aug 2012). This gene is co-expressed with the *USP18* gene in seven experiments, including GSE16147_d5, GSE13824_d1, and GSE6740_d22. The query result can be visualized as a graph representing the gene co-expression network as well as a tabular format.

**Figure 8:**
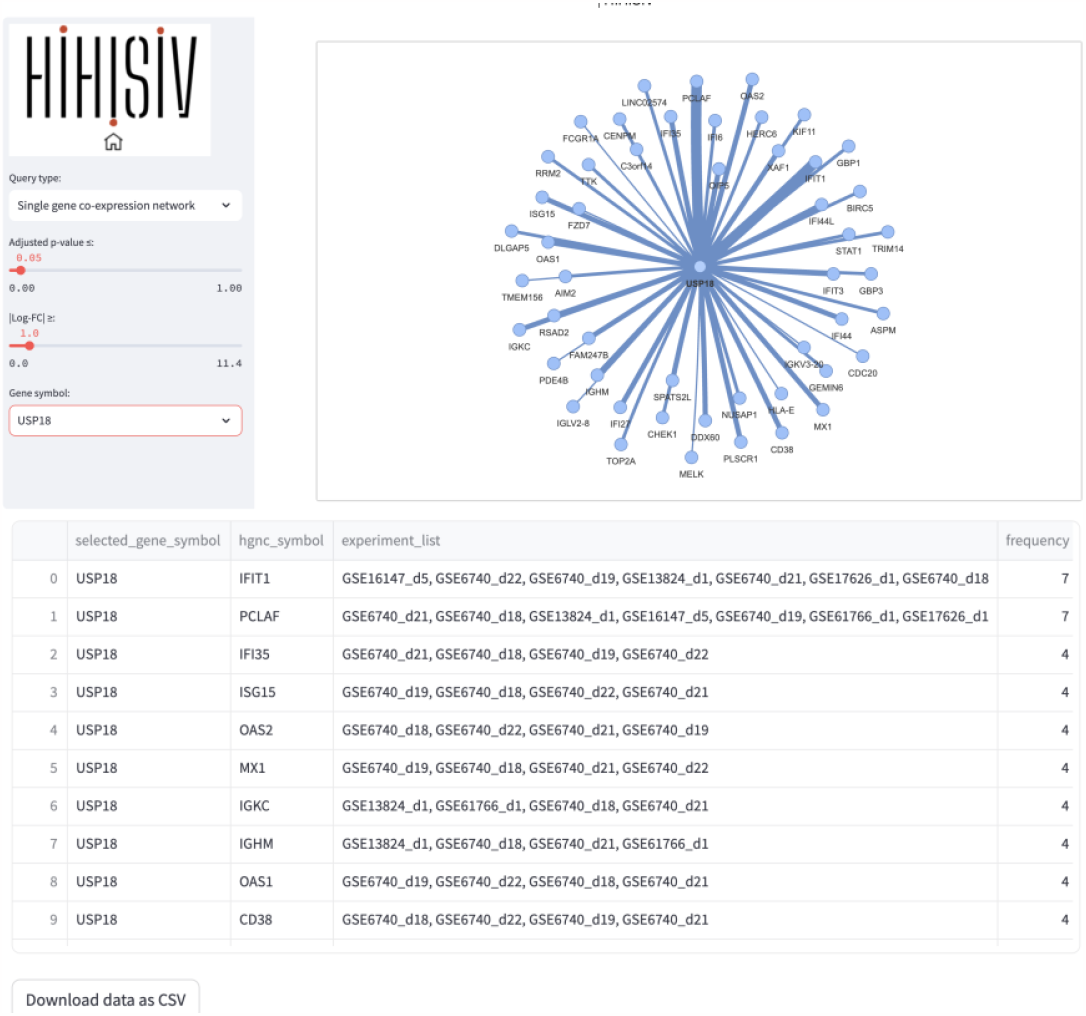
Example of ‘Single gene co-expressed network’ query for the gene *USP18*. The connections in this gene represent the genes co-expressed according to the parameters (adj_pvalue <= 0.05 and |log-FC| > 1) and the thickness of the connection varies according to the number of co-expressed experiments that the target gene has with the resulting gene.

#### Gene set co-expression network

In this query mode, instead of single gene co-expression, a set of genes can be entered and the result displays a visual network representation. The associated table facilitates a comprehensive understanding of the co-expression patterns and experimental associations among the genes of interest. For instance, in the example shown in Figure 9, we used the gene mentioned in the previous examples, *USP18*, along with other genes such as *STAT1, SP100, IFI35, APOBEC3A, MX1*, and *CXCR5* (adj_pvalue <= 0.05 and |log-FC| > 1). The query result reveals an interconnected network of genes associated with the immune response, particularly in defense against pathogens such as viruses. Additionally, the result is presented as a table indicating the experiments associated with each pair of connections. The user can also choose if disconnected nodes, i.e. genes that are not co-expressed with other genes, are also displayed in the network representation.

**Figure 9:**
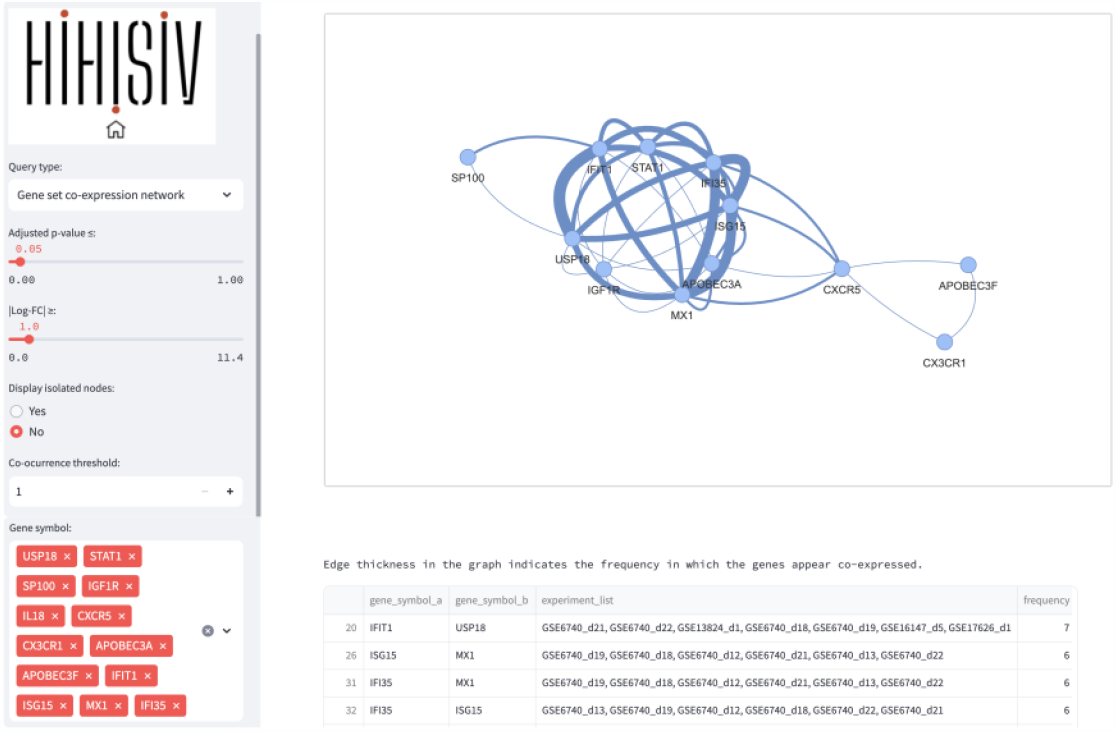
Result of the query module ‘Gene set co-expressed network’ (adj_pvalue <= 0.05 and |log-FC| > 1) shows an interconnected network of genes linked to the immune response, particularly in defense against pathogens such as viruses. The table result provides information on the experiments associated with each pair of connections.

## Conclusions

We present the HIHISIV database that provides a comprehensive integrated view of immune host response in SIV and HIV hosts. The data made available through HIHISIV, following best practices for supporting the FAIR principles, is based on aggregation of metadata and a workflow for analyzing Microarray and RNA-Seq datasets and annotations. The workflow identifies differentially expressed genes in the different studies analyzed and adds other types of interactions and relevant roles that these genes have. The HIHISIV database contains a web page with an easy-to-use interface for biologists to search and browse for genes and experimentally testable new hypotheses of molecular mechanisms related to the infection process in HIV/SIV and host types. The database also has additional information about viruses, documentation, experimental list and external sources. Our objective is to continue the development of the HIHISIV around the querying, metadata, analysis functionality, and addition of new datasets from main transcriptome repositories.

## Acknowledgements

The authors would like to thank the Brazilian National Laboratory for Scientific Computing for providing computational resources used in the analyses.

